# Oxytocin enhances excitability and potentiates synaptic transmission in dentate gyrus granule cells

**DOI:** 10.64898/2025.12.06.692683

**Authors:** Alyssa M. Marron, Brianna Balsamo, Darrin H Brager

## Abstract

The dentate gyrus is the principal gateway for information into the hippocampus. The dentate gyrus transforms input from the entorhinal cortex into sparse, selective representations that support memory formation. Oxytocin is a key neuromodulator of social behavior and supports social memory through its actions in hippocampal area CA2. However, whether oxytocin also modulates the dentate gyrus (DG), a source of major excitatory input to CA2, remains unclear. We performed whole-cell recordings to test if oxytocin modulated the excitability of mouse dentate gyrus granule cells. We found that bath application of the oxytocin receptor agonist Thy^4^, Gly^7^-oxytocin (TGOT) increased DG granule cell excitability by depolarizing the resting membrane potential, increasing input resistance, and hyperpolarizing action potential threshold. In addition to increasing postsynaptic excitability, we found that TGOT decreased the paired-pulse ratio of perforant path to granule cell synapses and also increased the frequency, without an effect on amplitude, of miniature EPSCs suggesting that TGOT increased the probability of glutamate release. Notably, long-term potentiation induced by theta-burst pairing occluded the effect of TGOT on synaptic strength suggesting that oxytocin and long-term potentiation may converge on common downstream mechanisms. Our results revealed a previously uncharacterized role for oxytocin in regulating the relay of information between the entorhinal cortex and dentate gyrus. This suggests a potential mechanism through which oxytocin shapes hippocampal processing of socially relevant stimuli.

**Significance Statement:** Oxytocin is essential for social memory and acts prominently within hippocampal area CA2, yet its influence on the dentate gyrus, a major excitatory input of CA2, has remained unclear. We demonstrate that oxytocin modulates dentate gyrus granule cell signaling by enhancing intrinsic excitability and strengthening perforant path synaptic transmission via increased presynaptic release probability. We further show that long-term potentiation occludes these synaptic effects suggesting convergence between oxytocin signaling and activity-dependent plasticity. These findings identify a previously unrecognized role for oxytocin in shaping dentate gyrus processing, broadening our understanding of how neuromodulatory signals influence hippocampal circuits involved in social information processing.

**Key points:**

- Oxytocin increases dentate gyrus granule cell excitability by depolarizing resting membrane potential, increasing input resistance and hyperpolarizing action potential threshold.
- Oxytocin enhances perforant path synaptic transmission by increasing presynaptic glutamate release probability.
- Long-term potentiation occludes the synaptic effects of oxytocin, suggesting overlap in downstream mechanisms.

## Introduction

Social memory, or the ability to recognize and remember conspecifics, is integral in learning to navigate the world as humans utilize social imitation to efficiently acquire new skills (Leblanc & Ramirez, 2020). The neuropeptide oxytocin, well-known for its roles in lactation, water retention, and childbirth (Cilz *et al*., 2018; Zagrean *et al*., 2022), is also important for social behaviors like pair bonding and emotional recognition (Dantzer *et al*., 1987; Young & Wang, 2004; Goldman & Marlow-O’Connor, 2009; Lee *et al*., 2009; Hurlemann & Grinevich, 2018). The hippocampus, and specifically area CA2, were previously implicated in social memory (Deng *et al*., 2010; Leuner *et al*., 2012; Cui *et al*., 2013; Piskorowski & Chevaleyre, 2013; Kohara *et al*., 2014; Wintzer *et al*., 2014; Hitti & Siegelbaum, 2014; Stevenson & Caldwell, 2014; Alexander *et al*., 2016; Dudek *et al*., 2016; Lin *et al*., 2017; Raam *et al*., 2017; Lin & Hsu, 2018; Cilz *et al*., 2018; Domínguez *et al*., 2019; Brown *et al*., 2020; Azahara *et al*., 2020; Zagrean *et al*., 2022; Lopez-Rojas *et al*., 2022; Loisy *et al*., 2022; Zhu *et al*., 2023). Oxytocin exerts its effects across various brain regions through activation of the G_q/11_ coupled receptor pathway (Gimpl & Fahrenholz, 2001). In CA2 pyramidal cells, this signaling cascade engages distinct branches of the G_q/11_ pathway to enhance intrinsic excitability and alter action potential shape (Tirko *et al*., 2018; Liu *et al*., 2022). Oxytocin also strengthens CA2 synaptic transmission through mechanisms requiring NMDA receptor activation, post synaptic calcium influx, and CaMKII signaling (Pagani *et al*., 2015), highlighting the capacity of oxytocin to modulate both intrinsic and synaptic properties within hippocampal circuits.

Pyramidal neurons in area CA2 receive strong, direct excitatory input from the dentate gyrus (DG) (Cui *et al*., 2013; Kohara *et al*., 2014; Hitti & Siegelbaum, 2014) and evidence suggests that the DG contributes to social memory processing. In rodents, DG granule cells (DG GCs) are strongly activated (indicated by high cFos expression) following a short-term social memory task (Bertoni *et al*., 2021), and disruption of the inputs from the entorhinal cortex (EC) to DG impairs social recognition (Leung *et al*., 2018). In the DG, oxytocin receptors were previously reported to be located in inhibitory interneurons and a small population of excitatory mossy cells (Mitre *et al*., 2016; Raam *et al*., 2017; Li *et al*., 2025) and were recently shown in the DG granule cell layer (Young & Song, 2020; Mitre *et al*., 2016). This suggests that oxytocin signaling in the DG may also play a role in social memory.

To our knowledge, there are no single cell electrophysiological studies on the effects of oxytocin on DG GCs. We therefore investigated the effects of the oxytocin receptor agonist Thy^4^, Gly^7^-oxytocin (TGOT) on DG GC function using whole-cell recordings in acute hippocampal slices. We found that bath application of TGOT increased input resistance, depolarized resting membrane potential, and hyperpolarized action potential threshold, resulting in increased DG GC excitability. We stimulated perforant path inputs to the DG and found that TGOT increased excitatory post synaptic potential (EPSP) slope and decreased paired pulse ratio, suggesting a presynaptic mechanism of oxytocin action. In agreement, TGOT increased the frequency, but not amplitude, of miniature excitatory postsynaptic currents (mEPSC). Finally, theta-burst pairing long-term potentiation occluded the effect of TGOT on synaptic transmission, indicating that oxytocin modulation may share common signaling pathways activity-dependent plasticity. Together, these results suggest a novel role of oxytocin in regulating DG GC excitability and modulation of hippocampal circuits involved in social memory processing.

## Methods

### Animals

All animal procedures were approved by the Institutional Animal Care and Use Committees at the University of Texas at Austin (AUP-2022-00257) or the University of Nevada, Las Vegas (IACUC-01217), and conformed to NIH guidelines. Acute hippocampal slices were prepared from 2–4 month-old C57BL/6J mice (JAX strain #000664) of both sexes. Sexes were pooled for all analyses. Mice were housed under a 12 h light/dark cycle with food and water available ad libitum.

### Slice preparation

Mice were anesthetized with a ketamine (100 mg/kg)/xylazine (10 mg/kg) cocktail and transcardially perfused with ice-cold saline containing (in mM): 2.5 KCl, 1.25 NaH_2_PO_4_, 25 NaHCO_3_, 0.5 CaCl_2_, 7 MgCl_2_, 7 dextrose, 205 sucrose, 1.3 ascorbate and 3 sodium pyruvate. The solution was continuously bubbled with 95% O_2_/5% CO_2_ to maintain a pH at ∼7.4. Coronal hippocampal slices (300 μm) were cut using a vibrating tissue slicer (Vibratome 3000, Vibratome Inc.). Slices recovered for 20 minutes at 35°C in a chamber filled with artificial cerebral spinal fluid (aCSF) containing (in mM): 125 NaCl, 2.5 KCl, 1.25 NaH_2_PO_4_, 25 NaHCO_3_, 2 CaCl_2_, 2 MgCl_2_, 10 dextrose and 3 sodium pyruvate (bubbled with 95% O_2_/5% CO_2_) and were then maintained at room temperature until the time of recording.

### Electrophysiology

Slices were transferred to a recording chamber and continuously perfused (1−2 mL/minute) with oxygenated aCSF containing (in mM): 125 NaCl, 3.0 KCl, 1.25 NaH2PO4, 25 NaHCO3, 2 CaCl2, 1 MgCl2, 10 dextrose, 3 sodium pyruvate (bubbled with 95% O_2_/5% CO_2_). The dentate gyrus was visualized using either a Zeiss Axioskop FS1 or AxioExaminer microscope. Whole-cell recordings were obtained from visually identified granule cells in the dentate gyrus of the hippocampus at 32–34°C or room temperature. Patch pipettes (3–8 MΩ) were filled with an internal solution containing (in mM): 120 K-gluconate, 16 KCl, 10 HEPES, 8 NaCl, 7 K_2_ phosphocreatine, 0.3 Na−GTP, 4 Mg−ATP (pH 7.3 with KOH). Neurobiotin (0.2%; Vector Laboratories) was included in the internal recording solution to confirm cell identity and recording location. Drugs were prepared from concentrated stock solutions in water and were obtained from Abcam pharmaceutical, Bachem or Tocris.

### Whole-cell recordings

Recordings were performed using a Dagan BVC−700 amplifier, ITC-18 (Instrutech) and AxoGraph X data acquisition software or a Sutter Integrated Patch Amplifier and SutterPatch data acquisition software. Data were acquired at 10−50 kHz and filtered at 5−10 kHz. Pipette capacitance was compensated and the bridge balance was adjusted before each recording. Series resistance was continuously monitored throughout each experiment and cells were excluded if series resistance exceeded 45 MΩ. Cells were held at −70 mV in current clamp unless otherwise noted.

For experiments measuring the intrinsic properties and excitability, ionotropic synaptic transmission was blocked by adding D-2-amino-5-phosphonopentanoate (D-AP5, 25 µM), 6,7-dinitroquinoxaline-2,3-dione (DNQX, 20 µM), and gabazine (GBZ, 2 µM) to the external solution. Firing rate was measured from responses to 500ms current injections (+50 to +500 pA, Δ50 pA). Input resistance was calculated from the linear portion of the current−voltage relationship generated from current steps (−50 to +50 pA, Δ10 pA). Action potential (AP) threshold was defined as the membrane potential at which dV/dt first exceeded 50 mV/ms.

### Synaptic Stimulation

A stimulating electrode was placed 200-400μm away from the recorded cell in the middle molecular layer, to activate the perforant path. The stimulus intensity was adjusted to evoke an initial excitatory post synaptic potential (EPSP) of ∼3-5mV. Pairs of stimuli (100 μs) were delivered at a 50 ms interstimulus interval and the paired-pulse ratio was calculated as the slope of EPSP2 divided by the slope of EPSP1.

Long-term potentiation was induced using a theta burst pairing (TBP) paradigm adapted from previously published protocols (Schmidt-Hieber *et al*., 2004; Kennedy *et al*., 2024). Briefly, EPSPs were evoked at 0.033Hz for 5 minutes followed by TBP. TBP consisted of ten staggered EPSP-AP pairs at 100Hz, delivered at 5Hz, and repeated four times at 10 sec intervals. Following TBP, EPSP slope was monitored at 0.033Hz for up to 30 minutes. In accordance with previously published TBP protocols in the DG (Schmidt-Hieber *et al*., 2004), dentate granule cells were held at -80mV during LTP experiments.

### mEPSC recordings

Miniature excitatory post synaptic currents (mEPSCs) were recorded in whole-cell voltage clamp at -80mV in the presence of 1 μM tetrodotoxin (TTX) and 2 μM GBZ. mEPSCs were collected in 500ms sweeps, repeated 360 times (total recording duration: 3 min.). Events were detected and analyzed using Sutterpatch 3.0’s synaptic event analysis program. The detection threshold was set to 1pA × 3 standard deviations, and a mEPSC template (2ms rise time, 3ms decay time) was used for event fitting. All detected synaptic events were manually filtered to exclude electrical noise.

### Post-hoc neuron visualization

At the end of recordings, slices were fixed in 3% glutaraldehyde at 4°C for a minimum of 24h. Slices were then washed with 0.1M phosphate buffer (6 × 5min), incubated in 1% Triton X-100-for 30 min, and subsequently incubated in 0.5% H_2_O_2_ for 30 minutes. Slices were washed in phosphate buffer (6 × 5min) and then incubated at 4°C for 24-72h in ABC reagent (Vector Laboratories) containing Avidin DH and biotinylated horseradish peroxidase. Slices were washed with 0.1M phosphate buffer (6 × 5min) and then incubated in DAB solution (Vector laboratories) in the presence of H_2_O_2_, resulting in a visible color change to the slices. Slices were dehydrated in glycerol and mounted on glass slides for imaging.

To determine the dorsal-ventral location of the hippocampal slices used for recordings, we mapped the dorsal ventral position of the histologically processed slices. Images for dorsal-ventral mapping were acquired using a Leitz Diaplan microscope at 5x magnification, and anatomical dimensions of hippocampal subfields were measured and scored according to previously published guidelines(Malik *et al*., 2016). Analysis showed that most cells were recorded from the mid-ventral region of the hippocampus (**Fig. 1A-B**).

**Figure 1.**
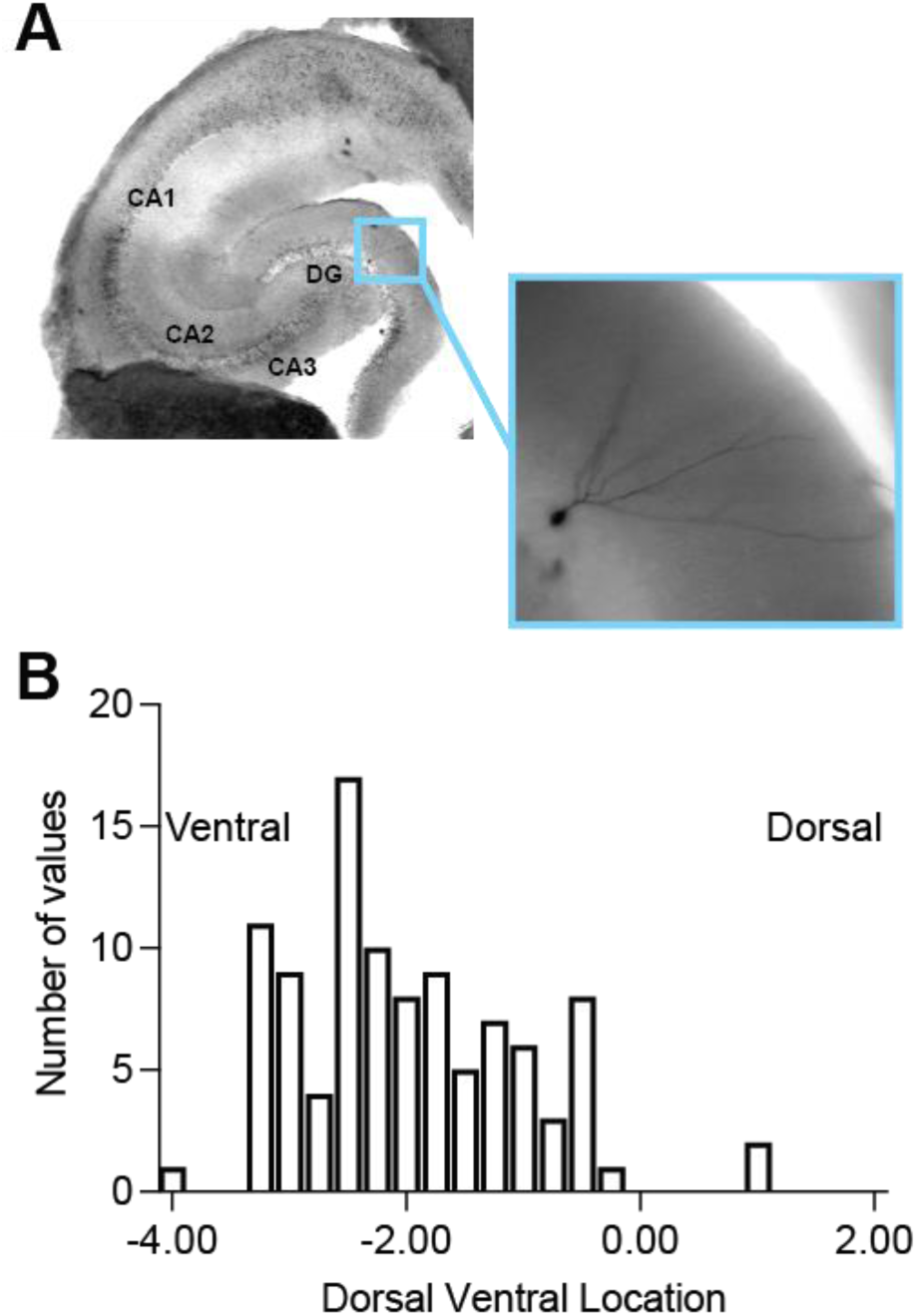
Dorsal ventral slice location of recorded dentate gyrus granule cells. *A*: representative DAB processed dentate granule cell showing granule cell layer location. *B*: dorsal-ventral location analysis, as measured in (Malik et al. 2016), showed recordings from dentate gyrus granule cells used in this study were in the mid-ventral region of the hippocampus.

### Data Analysis

Repeated-measures analysis of variance (ANOVA), mixed effects modules and post−hoc t−tests were used where appropriate to test for statistical differences between experimental conditions. Tukey corrections were used to correct for multiple comparisons. To quantify F–I curve shifts, the amount of current needed to elicit a firing rate of 20 Hz was interpolated from second-order polynomial curves (Tirko *et al*., 2018). Statistical analyses were performed in Prism (Graphpad). n refers to cells, with 1–4 cells recorded per mouse; m (number of animals) is reported in figure legends. Significance was set was set at α = 0.05 for all experiments.

## Results

### Oxytocin increases excitability of dentate granule cells

The oxytocin receptor agonist [Thr4,Gly7]-oxytocin (TGOT) increases the excitability and action potential firing rate of CA2 pyramidal neurons (Tirko *et al*., 2018). To investigate the effects of oxytocin on dentate gyrus granule cell excitability, we measured action potential firing in response to a series of depolarizing current injections before and after bath application of TGOT (400 nM). TGOT increased the firing rate of DG granule cells (**Fig. 2A-B**; F(1,22) = 6.393, n = 23, P = 0.0191, 2-way ANOVA). We quantified the increase in firing rate using two methods. First, we fit the input-output curve to a second order polynomial and quantified the current required to induce a 20Hz rate as previously described for the effect of TGOT on CA2 neurons(Tirko *et al*., 2018). We found a significant decrease in the current amplitude that elicited 20Hz firing (**Fig. 2C**; mean ± SEM, Baseline: 230.2 ± 17.81 pA, TGOT: 192.8 ± 17.05 pA, n = 20, P = <0.0001, Wilcoxon test). Second, we fit the linear portion of the F-I plot to measure the input-output gain(Hewitt *et al*., 2025). TGOT produced a small but significant increase in the neuronal gain of DG granule cells (**Fig. 2D**; Baseline: 0.09333 ± 0.005815 Hz/pA, TGOT: 0.09994 ± 0.005633 Hz/pA, n = 23, P = 0.0108, Wilcoxon test). Taken together, these results demonstrate that oxytocin robustly enhances the excitability of DG granule cells by lowering the current required to drive sustained firing and increasing input-output gain.

**Figure 2.**
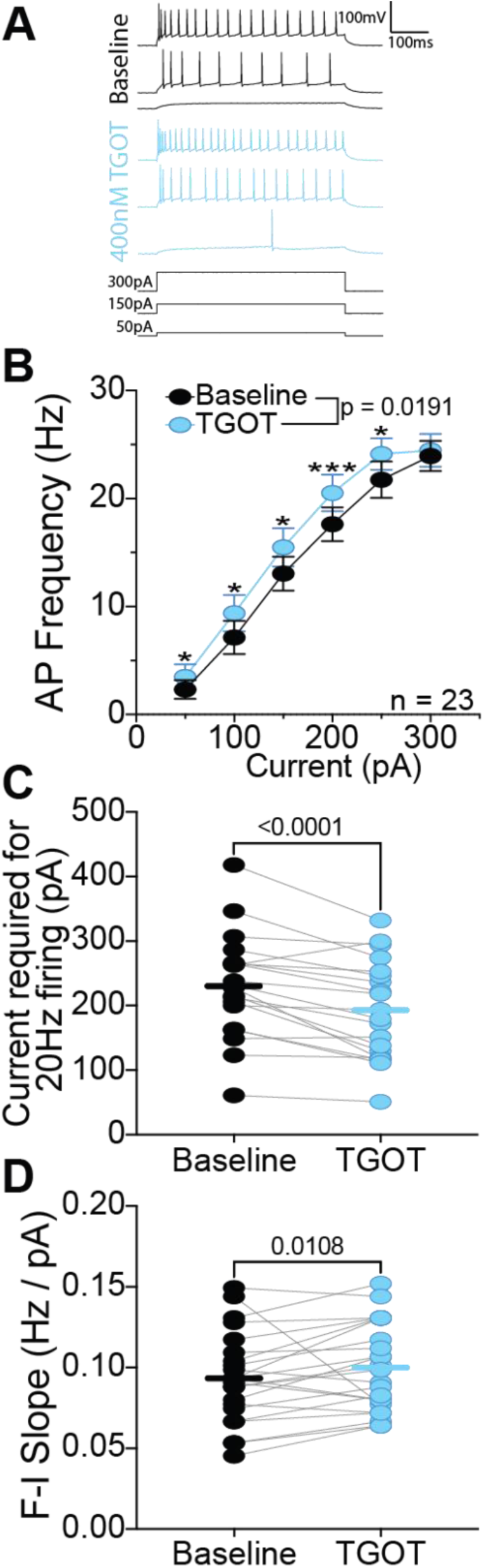
Oxytocin increases firing rate in dentate gyrus granule cells. A: representative voltage traces showing firing responses of DG granule cells to depolarizing current injections (bottom) before (top, black) and after bath application of TGOT (400 nM) middle, light blue). B: TGOT significantly increased the firing rate of DG granule cells (F(1,22) = 6.393, n = 23 cells, m = 13 mice, P = 0.0191, 2way ANOVA). C: to quantify the F-I curve shift, the input-output curve was fit to a second order polynomial and the current required to induce a 20Hz firing rate was interpolated as previously described in (Tirko et al. 2018). TGOT significantly reduced the current amplitude required to elicit 20 Hz firing (Baseline: 230.2 ± 17.81 pA; TGOT: 192.8 ± 17.05 pA; n = 20 cells, m = 12 mice, P < 0.0001, Wilcoxon test). D: TGOT significantly increased neuronal gain calculated from the linear range of the F–I curve (Baseline: 0.09333 ± 0.005815 Hz/pA; TGOT: 0.09994 ± 0.005633 Hz/pA; n = 23 cells, m = 13 mice, P = 0.0108, Wilcoxon test). Data are shown as mean ± SEM; n values are indicated in each panel.

### Oxytocin depolarizes the resting membrane potential and increases input resistance

Oxytocin depolarizes the resting potential and increases the input resistance of CA2 pyramidal neurons (Liu et al., 2022). We therefore asked if application of TGOT had similar effects on DG granule cells. We found that TGOT induced significant depolarization of resting membrane potential in DG granule cells (**Fig. 3A**; Baseline: -76.74 ± 2.035 mV, TGOT: - 73.89 ± 2.414 mV, n = 19, P = 0.0002, Wilcoxon test). Furthermore, TGOT significantly increased input resistance (**Fig. 3B-D**; F(1,22) = 9.728, P = 0.0050, mixed effects analysis) and alters the voltage-dependence of input resistance (**Fig. 3E**; Baseline: 5.705 ± 0.4784 MΩ/mV, TGOT: 4.708 ±0.5992 MΩ/mV, n = 20, P = 0.0362, Wilcoxon test). The more negative resting membrane potential of DG granule cells is largely driven by constitutive activity of G-protein coupled inwardly rectifying potassium channels (GIRK) (Gonzalez *et al*., 2018). In CA2 pyramidal neurons and neurons of the lateral nucleus of the central amygdala (CeL), oxytocin inhibits inwardly rectifying potassium channels (K_ir_) (Hu *et al*., 2020; Liu *et al*., 2022). We therefore tested if the depolarization of V_m_ and increase in R_N_ in DG granule cells by oxytocin were due to the inhibition of GIRK channels. Bath application of a low concentration of BaCl_2_ (25 µM), which blocks GIRK channels (Kim & Johnston, 2015; Malik & Johnston, 2017), significantly depolarized RMP (Baseline: -77.62 ± 0.8224 mV, low BaCl_2_: - 68.94 ± 2.251 mV, n = 8, P = 0.0156, Wilcoxon test). Subsequent application of TGOT, in the presence of BaCl_2_, further depolarized RMP (low BaCl_2_: - 68.94 ± 2.251 mV, TGOT: - 63.25 ± 2.608 mV, n = 8, P = 0.0156, Wilcoxon test) (F(1.566, 10.96) = 25.04, P = 0.0001, Rm one-way ANOVA) suggesting that in DG granule cells oxytocin depolarizes RMP independent of GIRK channels.

**Figure 3.**
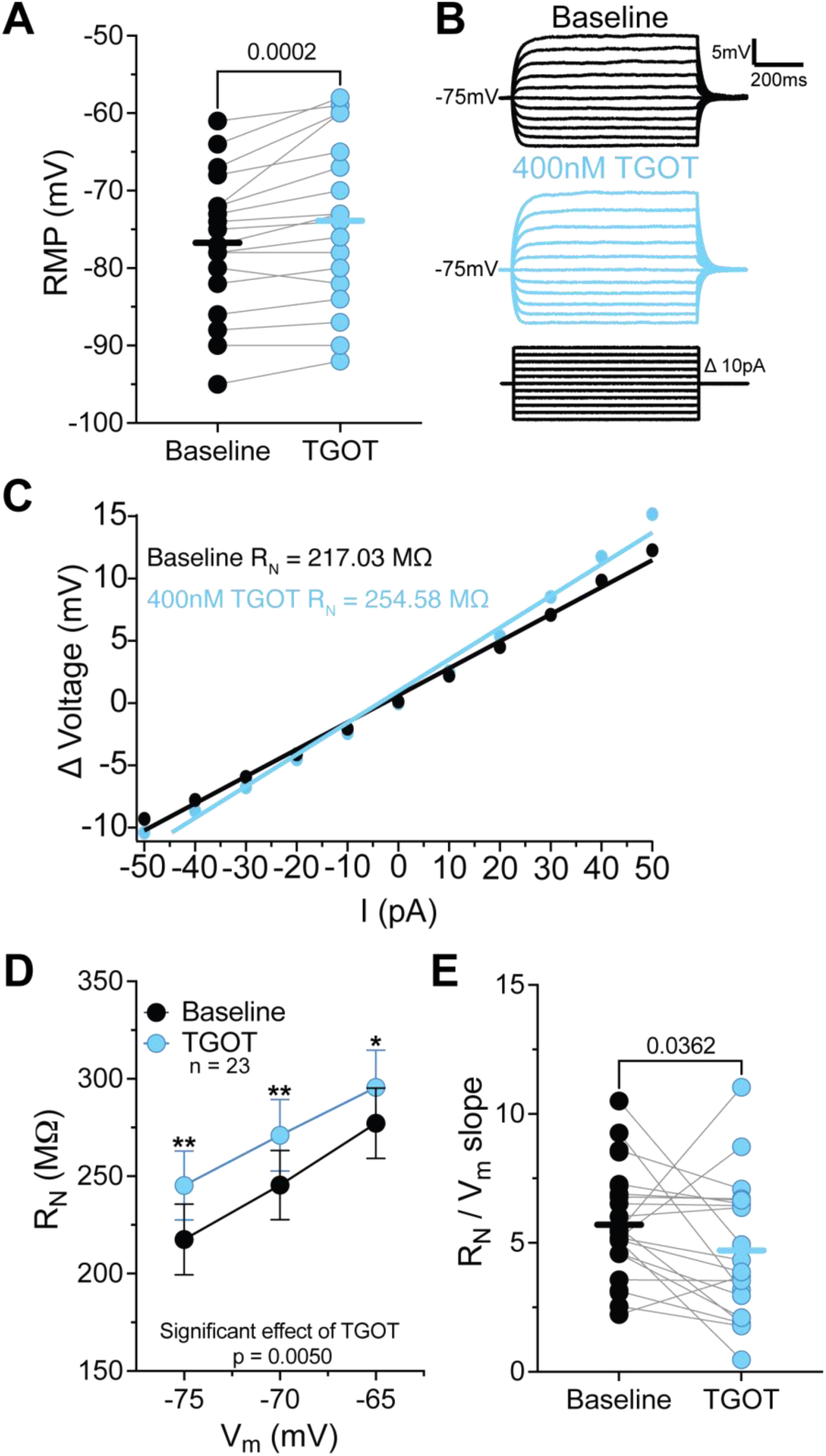
Oxytocin depolarizes RMP and increases input resistance in dentate gyrus granule cells. *A*: TGOT significantly depolarized resting membrane potential (RMP) (Baseline: –76.74 ± 2.04 mV; TGOT: –73.89 ± 2.41 mV; n = 19 cells, m = 9 mice, P = 0.0002, Wilcoxon test). *B*: representative voltage traces from DG granule cells in response to a series of current injections (*bottom*) before (*top, black*) and after bath application of TGOT (400 nM) (*middle, light blue*). *C*: representative current-voltage relationship showing input resistance calculation from the linear portion of the current−voltage relationship generated from current steps. *D*: input resistance significantly increased following TGOT application (F(1,22) = 9.728, n = 23 cells, m = 13 mice, P = 0.0050, mixed-effects analysis). *E*: voltage-dependence of input resistance was altered by TGOT (Baseline: 5.705 ± 0.4784 MΩ/mV; TGOT: 4.708 ± 0.5992 MΩ/mV; n = 20 cells, m = 12 mice, P = 0.0362, Wilcoxon test).

### Oxytocin hyperpolarizes action potential threshold by blocking K_V_1 channels

In addition to increasing input resistance, a change in the threshold of action potential generation would contribute to the increase in DG granule cell excitability by oxytocin. We used current steps of increasing duration and varied the amplitude to elicit a single action potential (**Fig. 4A-C**). Across all current durations, TGOT significantly hyperpolarized action potential threshold (**Fig. 4D**; F(1,22) = 14.08 n = 23, P = 0.0011, mixed effects analysis). TGOT had no effect on the minimum rate of change of voltage (min dV/dt) of a single AP (**Table 1**; F(1,22) = 1.606, n = 23, P = 0.2184, mixed effects analysis), but, interestingly, decreased the maximum rate of change of voltage (max dV/dt) (**Table 1**; F(1,22) = 5.892, n = 23, P = 0.0238, mixed effects analysis). Together, the results demonstrate that oxytocin increases DG granule cell excitability and neuronal gain by hyperpolarizing action potential threshold.

**Figure 4.**
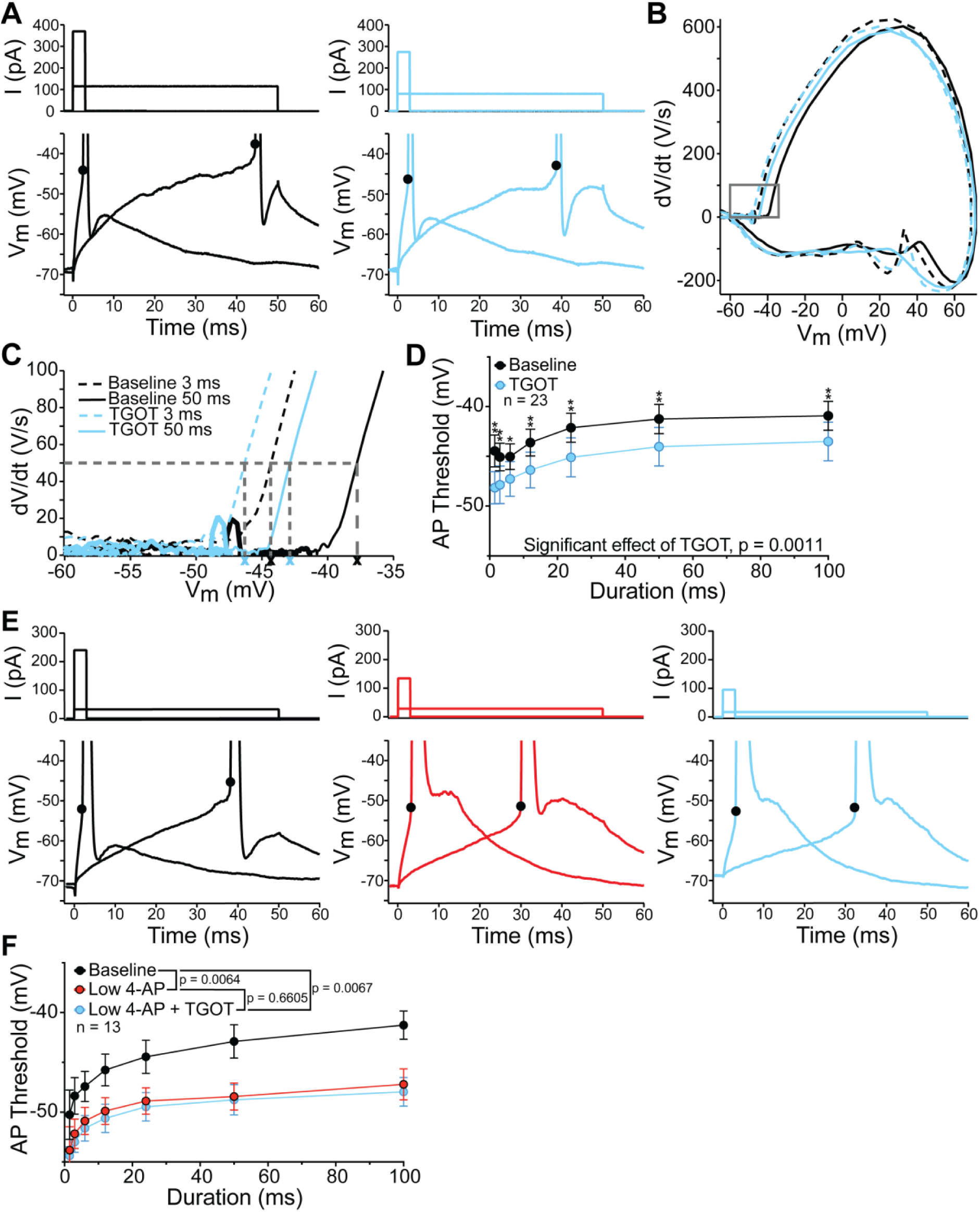
Oxytocin hyperpolarizes AP Threshold in dentate gyrus granule cells via modulation of K_V_1 channels. *A*: The action potential threshold (AP threshold) in DG granule cells was measured in response to current injections of increasing duration, that elicited a single action potential. Sample action potentials for short (*3ms*) and long (*50ms*) current injection durations before (*left, black*) and after bath application of TGOT (*right, light blue*). *B-C*: phase plane plot of action potentials in DG granule cells in response to 3ms and 50 ms current injections before (3ms, black dashed line; 50ms, black solid line) and after TGOT (3ms, light blue dashed line; 50ms, light blue solid line). AP threshold was defined as the membrane potential of the cell at which dV/dt first exceeded 50 mV/ms. *D*: TGOT significantly hyperpolarized AP threshold across all current durations (F(1,22) = 14.08, n = 23 cells, m = 13 mice, P = 0.0011, mixed-effects analysis). *E*: sample action potentials for short (*3ms*) and long (*50ms*) current injection durations at baseline (*left, black*), and after sequential bath application of low 4-AP (50 μM) (*middle, red*), followed by TGOT (400nM) (*right, light blue*). *F*: low 4-AP significantly hyperpolarized AP threshold (F(1,12)=10.83, P=0.0064) and occluded subsequent effects of TGOT (F(1,12)=0.2028, P=0.6605; n = 13 cells, m = 12 mice; mixed-effects analyses).

**Table 1.**
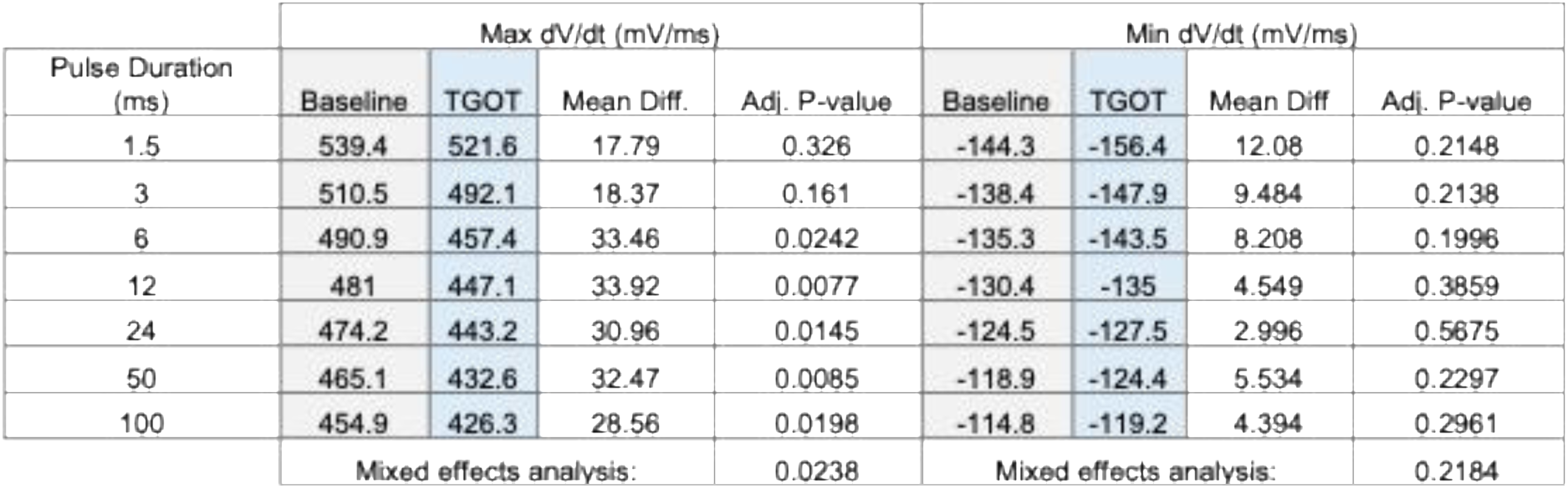
Oxytocin increased the maximum rate of change of action potentials. Maximum (max dV/dt) and minimum (min dV/dt) rates of change of membrane voltage during a single action potential were measured before and after bath application of TGOT (400 nM) in DG granule cells. TGOT significantly reduced max dV/dt (F(1,22) = 5.892, P = 0.0238, mixed effects analysis) but had no effect on min dV/dt (F(1,22) = 1.606, P = 0.2184, mixed effects analysis). Values are mean ± SEM; n = 23 cell, m = 13 mice.

| Pulse Duration<br>(ms) | Max dV/dt (mV/ms) |  |  |  | Min dV/dt (mV/ms) |  |  |  |
| --- | --- | --- | --- | --- | --- | --- | --- | --- |
|  | Baseline | TGOT | Mean Diff. | Adj. P-value | Baseline | TGOT | Mean Diff | Adj. P-value |
| 1.5 | 539.4 | 521.6 | 17.79 | 0.326 | -144.3 | -156.4 | 12.08 | 0.2148 |
| 3 | 510.5 | 492.1 | 18.37 | 0.161 | -138.4 | -147.9 | 9.484 | 0.2138 |
| 6 | 490.9 | 457.4 | 33.46 | 0.0242 | -135.3 | -143.5 | 8.208 | 0.1996 |
| 12 | 481 | 447.1 | 33.92 | 0.0077 | -130.4 | -135 | 4.549 | 0.3859 |
| 24 | 474.2 | 443.2 | 30.96 | 0.0145 | -124.5 | -127.5 | 2.996 | 0.5675 |
| 50 | 465.1 | 432.6 | 32.47 | 0.0085 | -118.9 | -124.4 | 5.534 | 0.2297 |
| 100 | 454.9 | 426.3 | 28.56 | 0.0198 | -114.8 | -119.2 | 4.394 | 0.2961 |
|  | Mixed effects analysis: |  |  | 0.0238 | Mixed effects analysis: |  |  | 0.2184 |

K_V_1 voltage-gated potassium channels significantly contribute to action potential threshold in multiple neuron types including DG granule cells (Morgan *et al*., 2019; Zbili *et al*., 2021; Lee *et al*., 2003). We next asked if oxytocin hyperpolarized action potential threshold by modulating K_V_1 channels. Bath application of low concentration 4-aminopyridine (50 µM 4-AP), which blocks K_V_1 channels, significantly hyperpolarized AP threshold (**Fig. 4E-F**; F(1,12) = 10.83, P = 0.0064, mixed effects analysis). Subsequent application of TGOT had no further effect on action potential threshold (**Fig. 4-F**; F(1,12) = 0.2028, P = 0.6605, mixed effects analysis). These data demonstrate that low 4-AP occludes the effect of TGOT on action potential threshold and suggests that oxytocin acts in part by blocking K_V_1 channels in DG granule cells.

### TGOT enhances presynaptic release probability at PP-DG granule cell synapses

Synaptic transmission from entorhinal cortex (EC) inputs to the dentate gyrus via the perforant path (PP) is critical for pattern separation and has been implicated in social recognition memory (Leung *et al*., 2018). However, whether oxytocin modulates DG granule cell synaptic transmission remains largely unknown. To determine if oxytocin modulates synaptic transmission at PP-DG granule cell synapses, we recorded pairs of excitatory postsynaptic potentials (EPSP) in response to stimulation of PP inputs in the middle molecular layer of the dentate gyrus. Bath application of TGOT significantly increased EPSP slope (**Fig. 5A-C**; Baseline: 0.5554 ± 0.04568 V/s, TGOT: 0.8125 ± 0.08980 V/s, n = 10, P = 0.0039, Wilcoxon test). The increase in EPSP slope was accompanied by a decrease in paired-pulse ratio after TGOT application (**Fig. 5D**; Baseline: 1.806 ± 0.1099, TGOT: 1.311 ± 0.1235, n = 10, P = 0.0059, Wilcoxon test), suggesting that the effect of TGOT had a presynaptic locus.

**Figure 5.**
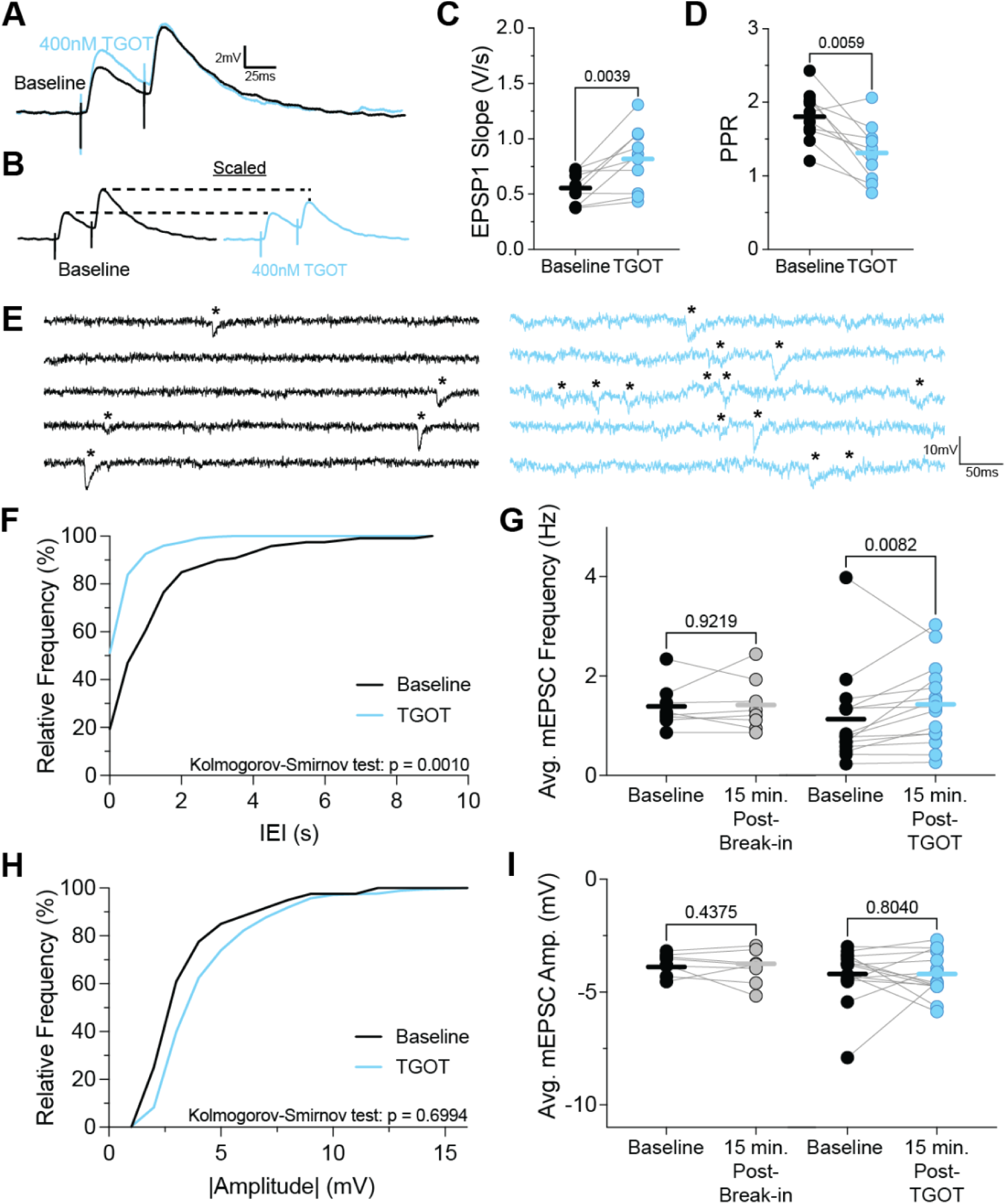
Oxytocin modulates PP-GC synapses through a presynaptic mechanism. *A*: averaged representative EPSPs recorded from DG granule cells in response to paired-pulse stimulation (interstimulus interval = 50 ms) of the middle molecular layer before (*black*) and after bath application of TGOT (*light blue*). *B*: scaled representative EPSPs from panel *A*. *C*: TGOT significantly increased EPSP slope (Baseline: 0.5554 ± 0.04568 V/s; TGOT: 0.8125 ± 0.08980 V/s, n = 10 cells, m = 7 mice, P = 0.0039, Wilcoxon test). *D*: paired-pulse ratio significantly decreased after bath application of TGOT (Baseline: 1.806 ± 0.110; TGOT: 1.311 ± 0.124, n = 10 cells, m = 7 mice, P = 0.0059, Wilcoxon test). *E*: representative traces from voltage-clamp recordings of mEPSCs (marked with *) from DG granule cells held at -80mV, in the presence of TTX and GBZ, before (*left, black*) and after bath application of TGOT (*right, light blue*). *F*: representative cumulative frequency distribution of mEPSC frequency (measured by interevent interval, IEI) before and after TGOT. TGOT significantly shifted the mEPSC IEI frequency distribution (D = 0.6316, P = 0.0010, Kolmogorov-Smirnov test). *G*: TGOT significantly increased average mEPSC frequency (*right; baseline, black; 15 min. post-TGOT, light blue*) (Baseline: 1.127 ± 0.238 Hz; TGOT: 1.425 ± 0.210 Hz; n = 15 cells, m = 9 mice, P = 0.0082, Wilcoxon test), with no effect observed in time-matched controls (*left; baseline, black; 15 min. post-break-in, grey*) (Baseline: 1.384 ± 0.1592 Hz, 15 minutes: 1.411 ± 0.1890 Hz, n = 8 cells, m = 3 mice, P = 0.9219, Wilcoxon test). *H*: representative cumulative frequency distribution of mEPSC |amplitude| before and after TGOT. TGOT significantly had no effect on the mEPSC |amplitude| frequency distribution (D = 0.2500, P = 0.6994, Kolmogorov-Smirnov test). *I*: TGOT did not change average mEPSC amplitude (Baseline: – 4.207 ± 0.311 mV; TGOT: –4.219 ± 0.246 mV; n = 15 cells, m = 9 mice, P = 0.8040), and amplitude was also unchanged in time-matched controls (Baseline: -3.749 ± 0.1677 mV, 15 minutes: -3.890 ± 0.2558 mV, n = 8 cells , m = 3 mice, P = 0.4375, Wilcoxon test).

To further test that the site of TGOT action was presynaptic, we recorded miniature excitatory post-synaptic currents (mEPSCs), in the presence of 1 μM tetrodotoxin (TTX) and 2 μM GBZ, before and after application of TGOT (**Fig. 5E**). Application of TGOT significantly increased mEPSC frequency (**Fig. 5F-G**; Baseline: 1.127 ± 0.2377 Hz, TGOT: 1.425 ± 0.2098 Hz, n = 15, P = 0.0082, Wilcoxon test) without any effect on mEPSC amplitude (**Fig. 5H-I**; Baseline: -4.207 ± 0.3110 mV, TGOT: -4.219 ± 0.2463 mV, n = 15, P = 0.8040, Wilcoxon test), consistent with an increase in presynaptic release probability. Time-matched controls (without TGOT application) showed no change in either mEPSC amplitude or frequency (**Fig. 5G,I**; Control mEPSC amplitude: Baseline: -3.749 ± 0.1677 mV, 15 minutes: -3.890 ± 0.2558 mV, n = 8, P = 0.4375, Wilcoxon test; Control mEPSC frequency: Baseline: 1.384 ± 0.1592 Hz, 15 minutes: 1.411 ± 0.1890 Hz, n = 8, P = 0.9219, Wilcoxon test). Together, these results suggest that oxytocin enhances PP–DG granule cell synaptic transmission by increasing the probability of glutamate release.

### LTP occludes the effect of oxytocin on synaptic transmission

Given that oxytocin appears to enhance glutamate release of PP synapses, we next asked what, if any, effect oxytocin had on long-term potentiation. We first wanted to assess the amount of potentiation that occurred using a previously published theta-burst pairing (TBP) protocol to induce LTP at PP-DG granule cell synapses (Schmidt-Hieber *et al*., 2004; Kennedy *et al*., 2024). In control experiments, TBP significantly increased the EPSP slope (**Fig. 6A-D**; Baseline: 1.144 ± 0.1083 mV/ms; TBP: 1.744 ± 0.3196 mV/ms, n = 9, P = 0.0195, Wilcoxon test), without a change in PPR (**Fig. 6E**; Baseline: 1.168 ± 0.1328, TBP: 1.101 ± 0.1178, P = 0.6523; n = 9, Wilcoxon test). These results are consistent with postsynaptic expression of LTP. In CA1 hippocampal neurons, activity-dependent changes in neuronal excitability (i.e., intrinsic plasticity) accompany TBP-LTP (Fan *et al*., 2005). To determine if intrinsic plasticity occurred after TBP in dentate gyrus granule cells, we measured the passive properties and action potential properties before and after TBP. We found LTP induction failed to increase or shift firing rate (**Fig. 6F**; F(1,8) = 2.286, P = 0.1690, 2way ANOVA) (**Fig. 6G**; Baseline: 231.3 ± 18.71 pA, TBP: 214.0 ± 16.91 pA, n = 9, P = 0.2500, Wilcoxon test), or increase R_N_ (**Fig. 6H**; Baseline: 170.1 ± 15.45 MΩ, TBP: 172.1 ± 13.11 MΩ, n = 9, P = 0.9102, Wilcoxon test). Following TBP-LTP, action potential threshold, in response to a 100 ms current step, was significantly hyperpolarized (**Fig. 6I-K**; Baseline: --38.02 ± 1.038 mV, TBP: -40.00 ± 1.227 mV, n = 11, P = 0.0137, Wilcoxon test). Taken together, these results suggest that TBP produces both synaptic potentiation and intrinsic plasticity in dentate gyrus granule cells.

**Figure 6.**
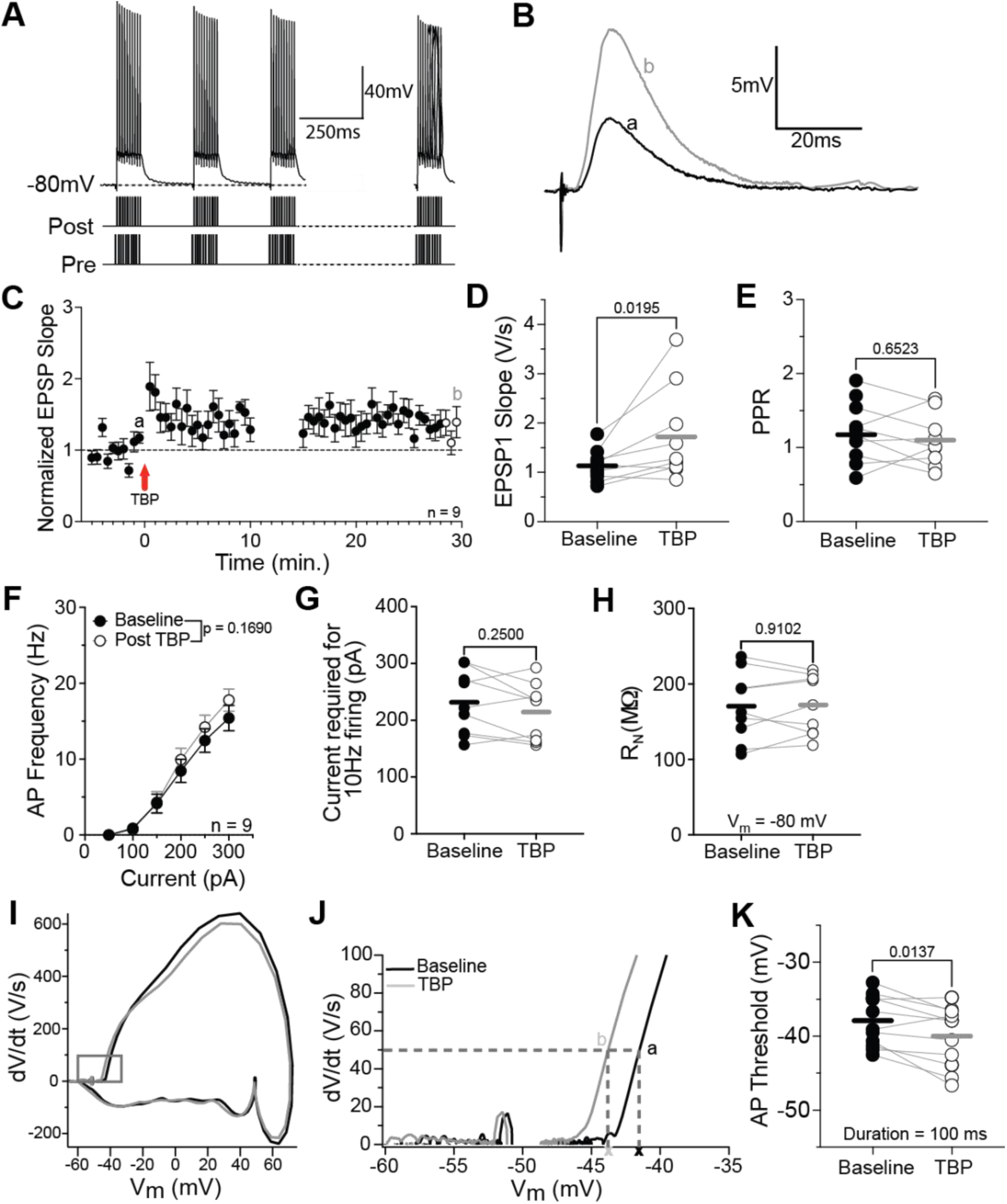
TBP induces LTP and hyperpolarizes AP threshold in dentate gyrus granule cells. *A*: Theta-burst pairing induction paradigms adapted from (Schmidt-Hieber et al. 2004). Cells were held at -80mV. *B*: averaged EPSPs recorded from DG granule cells in response to paired-pulse stimulation at time points indicated in panel *C*. C: Average plot of normalized EPSP slope in DG granule cells. *D*: TBP (*white*) significantly increased EPSP1 slope in DG granule cells compared to baseline (*black*) (Baseline: 1.144 ± 0.108 mV/ms; TBP: 1.744 ± 0.320 mV/ms, P = 0.0195, Wilcoxon test). *E*: PPR was unchanged after TBP (Baseline: 1.168 ± 0.1328, TBP: 1.101 ± 0.1178, P = 0.6523, Wilcoxon test). *F*: TBP did not alter DG granule cell firing rate (F(1,8) = 2.286, P = 0.1690, 2way ANOVA). *G*: the F-I curve shift was interpolated as described in Fig. 2 to calculate the current required to induce a 10Hz firing rate. TBP had no effect on the current required to elicit a 10Hz firing rate (Baseline: 231.3 ± 18.71 pA, TBP: 214.0 ± 16.91 pA, P = 0.2500, Wilcoxon test). *H*: TBP had no effect on input resistance (Baseline: 170.1 ± 15.45 MΩ; TBP: 172.1 ± 13.11 MΩ, P = 0.9102). *I-J*: phase plane plot of action potentials in DG granule cells in response to a 100 ms current injection before (black) and after TBP (grey). *K*: TBP significantly hyperpolarized AP threshold (Baseline: –38.02 ± 1.04 mV; TBP: –40.00 ± 1.23 mV, P = 0.0137, Wilcoxon test). *C-H*, *K*: n = 9 cells, m = 7 mice.

We then examined whether oxytocin enhances the potentiation of TBP induced-LTP in DG granule cells by having TGOT present throughout the recording, included at baseline (**Fig. 7A**). In the presence of TGOT, TBP increased EPSP slope (**Fig. 7B-D**; Baseline: 0.7966 ± 0.03121 mV/ms; TBP: 1.011 ± 1.07 mV/ms, n = 7, P = 0.0312, Wilcoxon test) without a change in PPR (**Fig. 7E**; Baseline: 1.340 ± 0.1188; TBP: 1.167 ± 0.1206, n = 7, P = 0.0781, Wilcoxon test), mirroring the effects of TBP in control LTP experiments. In the presence of TGOT, TBP continued to induce a significant hyperpolarization of AP threshold in response to a 100 ms current step (**Fig. 7F-H**; Baseline: -35.96 ± 1.019 mV, TBP: -38.39 ± 1.172 mV, n = 7, P = 0.0156, Wilcoxon test). These results indicate that oxytocin does not facilitate larger potentiation in DG granule cells. yet AP threshold hyperpolarizes after even in the presence of TGOT.

**Figure 7.**
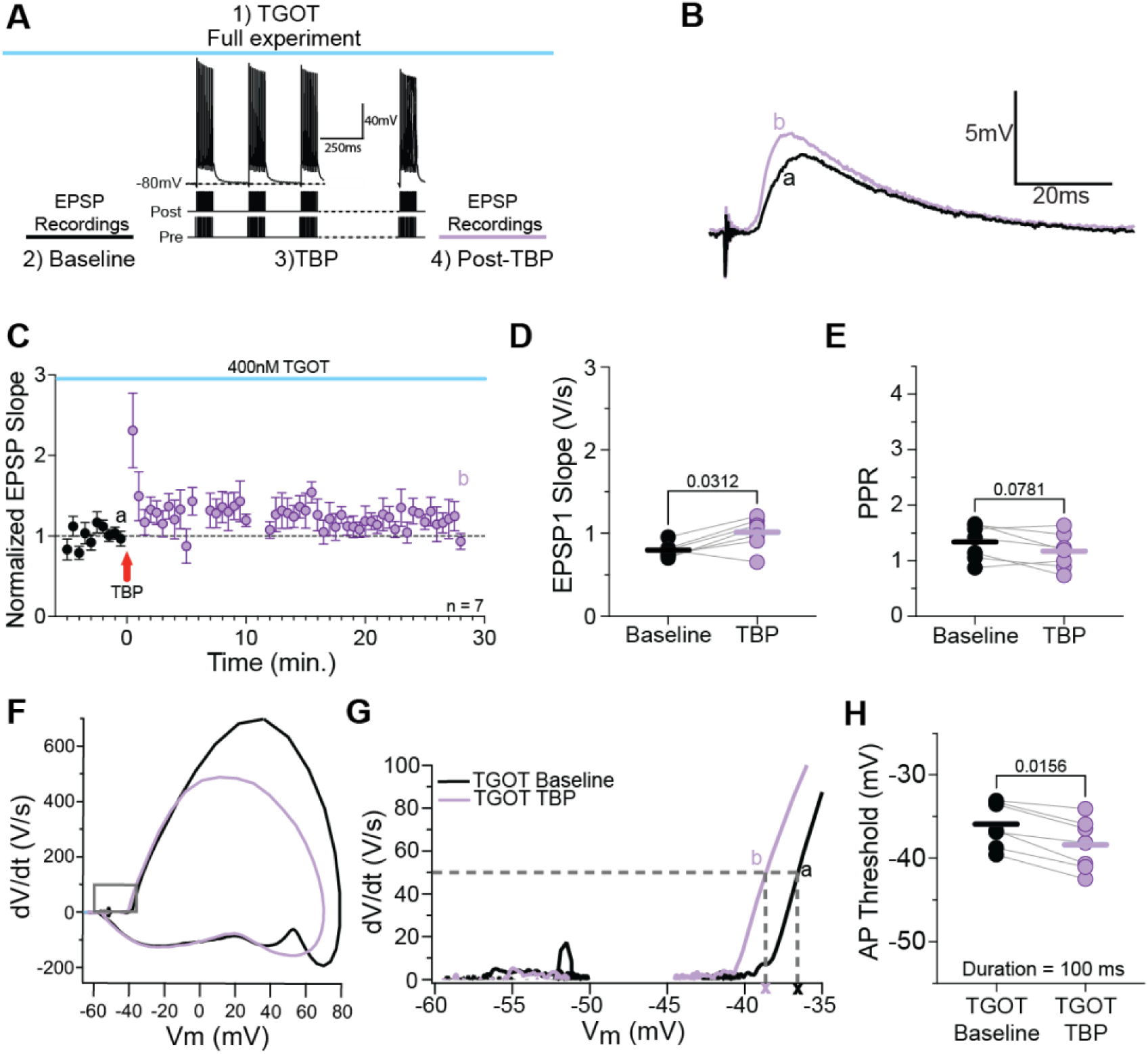
Oxytocin does not potentiate LTP-induced increase in EPSP slope. A: experimental timeline with TGOT (400nM) present throughout baseline (*black*) and LTP induction (*purple*). In order to account for TGOT-induced changes in RMP, cells were held at - 75mV. *B*: averaged EPSPs recorded from DG granule cells in response to paired-pulse stimulation at time points indicated in panel *C*. *C*: Average plot of normalized EPSP slope in DG granule cells. *D*: in the presence of TGOT, TBP produced a significant increase in EPSP slope (Baseline: 0.7966 ± 0.031 mV/ms; TBP: 1.011 ± 0.107 mV/ms, P = 0.0312, Wilcoxon test). *E*: in the presence of TGOT, TBP did not alter PPR (Baseline: 1.340 ± 0.1188; TBP: 1.167 ± 0.1206, P = 0.0781, Wilcoxon test). *F-G*: phase plane plot of action potentials in DG granule cells, in the presence of TGOT, in response to a 100 ms current injection before (*black*) and after TBP (*purple*). *H*: TBP hyperpolarized AP threshold despite continuous TGOT (Baseline: –35.96 ± 1.02 mV; TBP: –38.39 ± 1.17 mV, P = 0.0156, Wilcoxon). *C-E*, *H*: n = 7 cells, m = 4 mice.

We next asked if oxytocin could potentiate synaptic transmission after LTP induction. First, LTP was induced by TBP, and after a 5-minute post-TBP period, TGOT was applied for the remainder of the recording (**Fig. 8A**). As expected, TBP significantly increased EPSP slope (**Fig. 8B-D**). Surprisingly however, TGOT had no additional effect on EPSP slope (**Fig. 8D**; F(1.799,14.39) = 6.939, P = 0.0091; Baseline: 1.1 ± 0.14 mV/ms, TBP: 1.7 ± 0.19 mV/ms, TGOT: 1.5 ± 0.22 mV/ms, RM one-way ANOVA). TGOT also had no effect on PPR after TBP (**Fig. 8E**; F(1.196,8.566) = 0.7990, P = 0.4157; Baseline: 1.5 ± 0.25, TBP: 1.4 ± 0.19, TGOT: 1.6 ± 0.15, RM one-way ANOVA). These findings suggest that while oxytocin enhances glutamate release under baseline conditions, the effect is occluded after LTP induction, suggesting that oxytocin may engage mechanisms with both pre- and postsynaptic effects to alter synaptic transmission.

**Figure 8:**
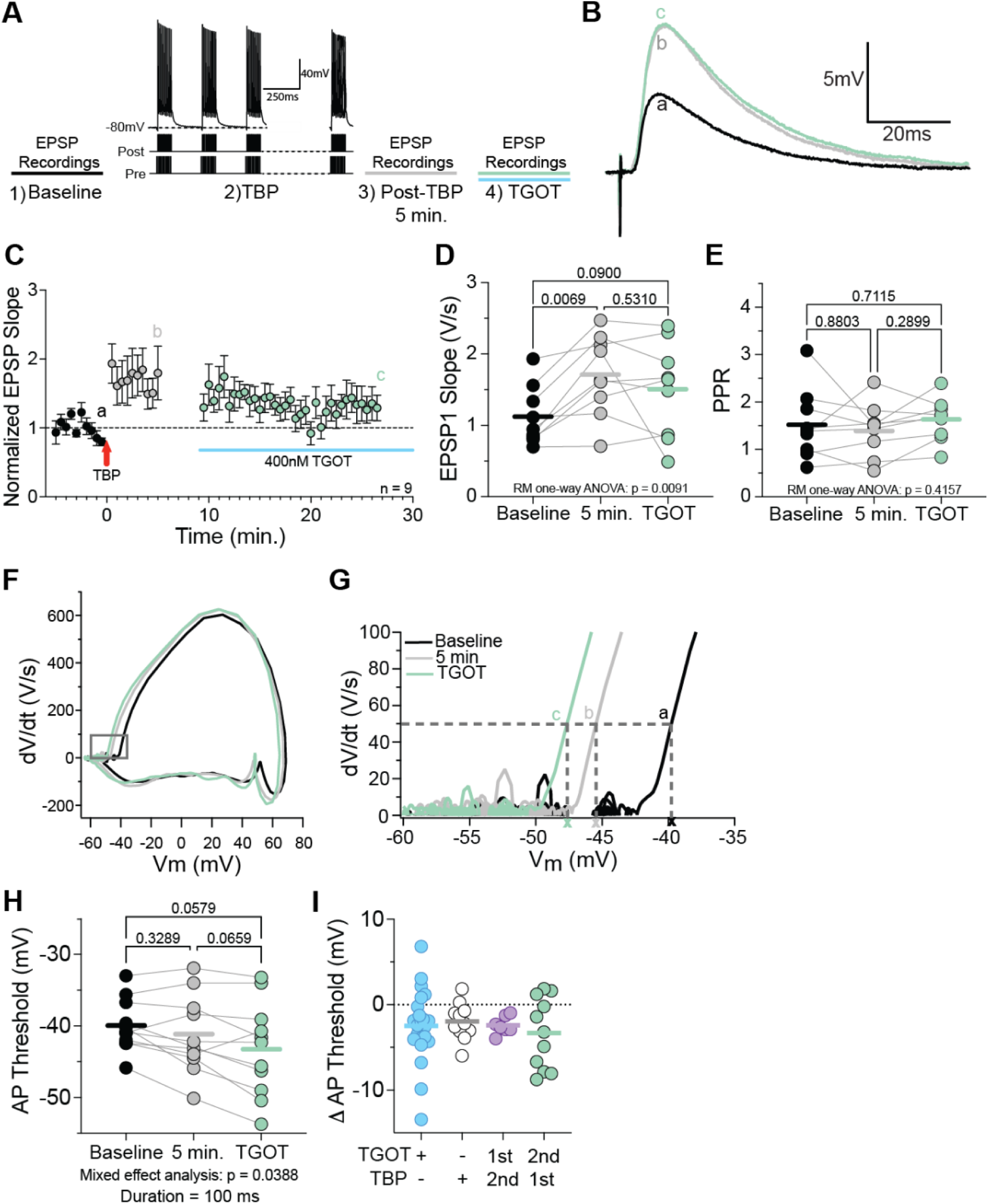
LTP Induction occludes effect of oxytocin on EPSP slope and AP threshold. *A*: experimental timeline illustrating 5 minutes baseline EPSP recordings (*black, a*), TBP delivery to DG granule cells (2), followed by a 5-minute EPSP recording period (*grey, b*) and a subsequent 15-minute bath application of TGOT (*green, c*). Cells were held at -80mV. *B*: averaged EPSPs recorded from DG granule cells in response to paired-pulse stimulation at time points indicated in panel *C*. *C*: Average plot of normalized EPSP slope in DG granule cells (n = 9 cells, m = 4 mice). *D*: TBP increased EPSP slope, but TGOT produced no additional potentiation (F(1.799,14.39) = 6.939, P = 0.0091; Baseline: 1.1 ± 0.14 mV/ms; TBP: 1.7 ± 0.19; TGOT: 1.5 ± 0.22, RM one-way ANOVA). *E*: PPR was unchanged across conditions (F(1.196,8.566) = 0.7990, P = 0.4157; Baseline: 1.5 ± 0.25; TBP: 1.4 ± 0.19; TGOT: 1.6 ± 0.15, RM one-way ANOVA). *F-G*: phase plane plot of action potentials in DG granule cells in response to a 100 ms current injection at baseline (*black*), 5 minutes post-TBP (*grey*), and following subsequent application of TGOT (*green*). *H*: Following TBP, TGOT did not further hyperpolarize AP threshold (F(0.9659,9.176) = 0.4830, P = 0.0388; Baseline: – 39.96 ± 1.09 mV; 5 min post-TBP: –41.16 ± 1.78; TGOT: –43.27 ± 1.96, RM one-way ANOVA). *I*: All experimental conditions, TGOT (*blue*, from Fig. 4D), TBP (*white*, from Fig. 6K), TBP in the presence of TGOT (*purple*, from Fig. 7H), or TBP followed be sequential application of TGOT (*green*, from Fig. 8H) produced similar magnitudes of AP threshold hyperpolarization (TGOT: - 2.466 ± 0.9062 mV, TBP: -1.970 ± 0.6411 mV, TGOT followed by TBP: -2.423 ± 0.3895 mV, TBP followed by TGOT: -3.311 ± 1.249 mV, P = 0.8845, Kruskal-Wallis test). *C-E*, *H*: n = 9 cells, m = 4 mice.

Following LTP induction, subsequent application of TGOT did not produce an additional significant hyperpolarization of AP threshold (**Fig. 8F-H**; F(0.9659,9.176) = 0.4830, P = 0.0388; Baseline: -39.96 ± 1.089 mV, 5 min.: -41.16 ± 1.777, TGOT: -43.27 ± 1.961 mV, RM one-way ANOVA), indicating that LTP induction occludes the effect of oxytocin on AP threshold. Finally, we assessed whether oxytocin application, TBP alone, TBP delivered in the presence of TGOT, or sequential TBP followed by TGOT differentially hyperpolarized AP threshold. Across all conditions, AP threshold was hyperpolarized to a similar degree (**Fig. 8I**; Δ AP Threshold: TGOT: -2.466 ± 0.9062 mV, TBP: -1.970 ± 0.6411 mV, TGOT followed by TBP: -2.423 ± 0.3895 mV, TBP followed by TGOT: -3.311 ± 1.249 mV, P = 0.8845, Kruskal-Wallis test).

## Discussion

Although the importance of oxytocin signaling in hippocampal area CA2 to social memory is well established, whether oxytocin plays a role in other hippocampal regions is not known. We found granule cells in the dentate gyrus are modulated by oxytocin. We observed that the oxytocin receptor agonist TGOT increases dentate gyrus granule firing rate, input-output gain, input resistance, and hyperpolarized action potential threshold. Together these results suggest that oxytocin increases the excitability of dentate gyrus granule cells. In parallel, TGOT increased the frequency of observing mEPSCs with no effect on mEPSC amplitude. In addition, TGOT decreased the paired-pulse ratio of EPSP evoked by stimulation of inputs in the middle molecular layer of the dentate gyrus. These observations suggest that oxytocin increases the probability of glutamate release from perforant path to granule cell synapses. We asked how these effects of oxytocin may affect activity-dependent plasticity in dentate granule cells. Theta-burst pairing potentiated perforant path inputs to granule cells. Interestingly, TBP also produced plasticity of intrinsic excitability which was observed as a hyperpolarization of action potential threshold. The interaction between oxytocin and long-term plasticity yielded confounding observations. First, TBP in the presence of oxytocin produced equivalent LTP as observed under control conditions. When oxytocin was applied after TBP however, we did not see any further synaptic potentiation. These results have interesting implications for how the dentate gyrus may encode socially relevant stimuli.

### Oxytocin enhances DG GC excitability via mechanisms paralleling other hippocampal subfields

Our data reveal that oxytocin robustly enhances DG GC excitability through increased firing rate, F-I slope, and R_N_, depolarization of RMP, and hyperpolarization of AP threshold. These effects are consistent with previously published results on the effect oxytocin in CA2 pyramidal neurons (Tirko *et al*., 2018; Liu *et al*., 2022). In CA2 pyramidal neurons, oxytocin receptors are proposed to modulate excitability via a G_q/11_–PLC–dependent signaling pathway that reduces M-current potassium channel (M-channels) activity. In DG granule cells, M-channels similarly influence excitability and somatic R_N_ (Mateos-Aparicio *et al*., 2014). Blocking these channels in DG granule cells increases firing rate, F-I slope, and R_N_, effects that parallel those observed following TGOT application and highlight M-channels as a potential target of oxytocin signaling in this population (Mateos-Aparicio *et al*., 2014). However, prior work in CA2 pyramidal neurons reported that oxytocin receptor activation predominantly suppressed an inward current mediated by inhibition of inwardly rectifying potassium channels (K_ir_), but not M-channels, suggesting that oxytocin may modulate excitability through multiple potassium conductances (Liu *et al*., 2022). Consistent with this hypothesis, oxytocin reduces K_ir_ activity via inhibition of GIRK channels in neurons of the central amygdala (Hu *et al*., 2020). We found that block of GIRK channels, with a low concentration of barium, did not occlude the effects of oxytocin. While this suggests that oxytocin does not modulate GIRK channels in DG granule cells, these findings do not rule out the possibility that oxytocin may block other inwardly rectifying potassium channels.

One notable difference between prior CA2 studies and our investigation of DG granule cells, is the hyperpolarization of action potential threshold by oxytocin. In CA2 neurons, oxytocin did not affect action potential threshold (but did produce a depolarization of the membrane potential and increase input resistance) (Tirko *et al*., 2018). We hypothesized that the hyperpolarization of threshold by oxytocin in DG granule cells is due to inhibition of voltage-gated K^+^ channels which participate in the setting of action potential threshold. K_V_1 channels are well-suited to modulate action potential threshold because they are closed at rest, activate with small depolarizations near subthreshold potentials, and produce outward currents that can control excitability (D’Adamo *et al*., 2020). We found that block of K_V_1 channels with low 4-AP occluded the hyperpolarization of threshold by oxytocin. This implicates K_V_1 channels as downstream targets of oxytocin signaling and positions them alongside M-channels and K_ir_ currents as candidate ion channels through which oxytocin may tune excitability in DG granule cell neurons. These results support the emerging view that oxytocin engages distinct potassium channel families across cell types, and suggest that in DG granule cells, modulation of K_V_1 channels represents a mechanism by which oxytocin receptor activation sharpens spike responsiveness and enhances the integrative properties of this sparsely firing population. Further characterizing the potassium currents modulated by oxytocin will be an important next step toward understanding the mechanisms by which oxytocin enhances intrinsic DG GC excitability. Together, these results indicate that oxytocin directly alters DG excitability through convergent modulation of subthreshold membrane properties, a mechanism that may tune the input–output transformation of the PP–DG circuit.

### Synaptic effects and presynaptic modulation in the DG

The DG integrates overlapping synaptic input from the EC and transforms DG GC firing in response to subtle environmental changes. This process, known as pattern separation, allows the activation of distinct DG GC populations to encode different contextual information (Leutgeb *et al*., 2007; Bakker *et al*., 2008; Deng *et al*., 2013). Pattern separation is critical for certain forms of recognition memory, specifically, recollection of events with shared features (Yassa & Stark, 2011). While the DG’s role in pattern separation is well-defined, its contribution to social memory remains less clear, and it is unknown if pattern separating social inputs is a distinct process from encoding other types of stimuli.

Our findings indicate that, in parallel with the effects on DG granule cell excitability, oxytocin signaling can enhance synaptic transmission at perforant path to granule cell synapses. Bath application of TGOT increased the slope of EPSPs and decreased the paired-pulse ratio (PPR) at PP-DG granule cell synapses. Furthermore, oxytocin increased mEPSC frequency, without affecting amplitude, supporting a presynaptic increase in glutamate release probability at PP-DG synapses. Importantly, these presynaptic effects persisted in the presence of TTX, indicating that oxytocin modulates presynaptic terminals independently of network activity or upstream neuronal firing. Similar presynaptic effects were observed in the auditory and piriform cortices, where oxytocin increases spontaneous event frequency through modulation of inhibitory terminals (Mitre *et al*., 2016). Together, these results suggest that oxytocin can amplify entorhinal input to the DG, potentially sharpening the activation of granule cell ensembles and enhancing the network’s ability to perform pattern separation in behaviorally relevant contexts.

### Oxytocin shares some signaling pathways with LTP

In addition to its effects on basal excitability and presynaptic transmission, oxytocin signaling interacts with activity-dependent synaptic plasticity in DG granule cells. In our study, theta-burst pairing (TBP) LTP occluded TGOT-induced enhancement of EPSP slope, indicating that the mechanisms recruited by oxytocin and by classical LTP partially overlap. DG granule cells receive synaptic input from both the medial and lateral entorhinal cortex (MEC and LEC), and LTP at these pathways can arise through both postsynaptic and presynaptic contributions. MEC-DG LTP is generally considered post synaptic (Christie & Abraham, 1994), though presynaptic contribution were reported in the guinea pig hippocampus (Min *et al*., 1998). In contrast, LEC-DG LTP increases presynaptic release probability (Christie & Abraham, 1994), mirroring the presynaptic enhancement we observe with oxytocin. This form of LTP is hypothesized to involve a postsynaptic NMDAR-dependent retrograde signal that modulates presynaptic terminals (Christie & Abraham, 1994).

Paired-pulse stimulation of these pathways further support this distinction: MEC inputs generally produce paired-pulse depression (PPR<1), whereas LEC inputs show paired-pulse facilitation (PPR>1) (Christie & Abraham, 1994). Our initial oxytocin paired-pulse experiments yielded PPRs greater than 1, suggesting preferential recruitment of LEC inputs. However, spontaneous mEPSCs reflect transmission from both MEC and LEC inputs, and PPR values across all LTP experiments showed both facilitation and depression. Thus, our stimulation likely activated a mixture of MEC and LEC inputs, limiting our ability to isolate pathway-specific oxytocin effects.

Despite this complexity, the findings that TBP induced LTP occludes oxytocin’s enhancement of EPSP slope suggests that oxytocin and LTP ultimately converge onto shared downstream pathways, potentially engaging overlapping NMDAR-dependent mechanisms. This interpretation is consistent with findings from CA1 and CA2, where oxytocin similarly potentiates synaptic transmission and recruits LTP-related signaling cascades. In CA1, oxytocin potentiates synaptic transmission and promotes the transition from early- to late-phase LTP through MAP kinase activation, CREB phosphorylation, and protein synthesis, without affecting LTP induction or basal transmission (Tomizawa *et al*., 2003; Lin *et al*., 2012). In CA2, oxytocin drives potentiation of EC to CA2 synapses via NMDAR- and CaMKII-dependent mechanisms NMDAR-and CaMKII-dependent mechanisms (Pagani *et al*., 2015; Lin *et al*., 2018; Tirko *et al*., 2018).

Clarifying how presynaptic oxytocin signaling intersects with postsynaptic LTP mechanisms in DG granule cells will be an important direction for future work.

Notably, TBP also hyperpolarized the action potential threshold, mirroring one of the postsynaptic effects of TGOT. This TBP-induced shift in AP threshold persisted even when oxytocin signaling was already engaged, and TGOT failed to produce any additional hyperpolarization when applied after TBP. Interestingly, despite producing a hyperpolarization of AP threshold comparable to that induced by oxytocin, TBP did not alter firing rate, neuronal gain, or input resistance. One possibility is that TBP recruits a mechanism that shift threshold without strongly influencing the currents that determine firing during sustained depolarization, whereas oxytocin engages a broader set of conductances that boost overall excitability. Thus, even though both manipulations shift AP threshold, the downstream effects on excitability may diverge depending on the specific conductances each pathway engages.

It is also notable that AP threshold measured at 5 minutes post-induction was not yet significantly hyperpolarized, raising the possibility that LTP dependent changed in AP properties may develop on a slower timescale than immediately detectable post-induction. Since our experiments were not designed to dissect the underlying channel mechanisms, we cannot determine whether oxytocin and LTP recruit overlapping or distinct mechanisms to converge at the level of spike initiation.

From this perspective, oxytocin increases DG granule cell excitability and enhances PP-DG synaptic transmission under baseline conditions, but these effects are not expressed once synaptic strength has already been elevated by LTP. Instead, oxytocin and LTP may independently modulate granule cell spike initiation through distinct mechanisms that nevertheless converge on similar functional outcomes. By engaging separate but complementary pathways, oxytocin and LTP may allow the DG to dynamically regulate excitability, with oxytocin providing state-dependent modulation and LTP enabling activity drive, experience-dependent adjustments.

### Circuit-level implications of oxytocin signaling in the dentate gyrus

Our study focused exclusively on excitatory signaling in the DG, leaving the effects of oxytocin on local inhibitory circuits unexplored. Nevertheless, oxytocin’s actions in the DG likely also reshape the local balance between excitation and inhibition. Previous work shows that oxytocin depolarizes fast-spiking hilar interneurons, increasing GABA release onto mossy cells (Harden & Frazier, 2016), while in CA2, it enhances both excitatory and inhibitory neuron excitability but ultimately biases the circuit toward net excitation (Tirko *et al*., 2018). Our findings indicate that oxytocin directly modulates presynaptic terminals to enhance glutamate release onto DG granule cells, providing a mechanism for dynamically tuning excitatory input to the hippocampus. These findings of increased DG GC excitability and glutamate release probability are consistent with a similar shift in the excitation–inhibition ratio to area CA2, potentially enhancing the transmission entorhinal cortex inputs. Such modulation could refine population coding within the granule cell layer, facilitating sparse but reliable pattern separation, a fundamental computational function of the DG.

By enhancing excitatory transmission and lowering AP threshold, oxytocin may also facilitate the induction of LTP by subthreshold stimulation, similar to its effects in CA1, where it lowers the threshold for protein synthesis-dependent long term-LTP (Lin *et al*., 2012). Together, these findings suggest that oxytocin acts at multiple sites within the DG circuit, increasing granule cell excitability via postsynaptic mechanisms while simultaneously enhancing presynaptic release probability at perforant path synapses. This positions oxytocin as a key regulator of both cellular and synaptic determinants of DG function, with presynaptic enhancement potentially representing an early permissive step for long-lasting potentiation.

### Behavioral and functional relevance

At the behavioral level, oxytocin signaling within the DG may contribute to social memory persistence and contextual encoding. Deletion of oxytocin receptors in the anterior hilus impairs social recognition (Raam *et al*., 2017), and activation of dorsal DG GCs correlates with social interaction (Bertoni *et al*., 2021). Our physiological findings further show that oxytocin enhances DG GC excitability and potentiates synaptic transmission, paralleling its facilitation of excitability and transmission in CA1 and CA2. Therefore, it is possible that oxytocin may promote the consolidation of social hippocampal-dependent memories by coordinating excitatory throughput and intrinsic excitability across DG and downstream brain regions. Distributed modulation of this kind could underlie oxytocin’s ability to facilitate social memory encoding, and future in vivo experiments with circuit-specific manipulations will be critical for clarifying how DG oxytocin signaling integrates into broader hippocampal networks to regulate social cognition. Overall, oxytocin signaling provides a mechanism for amplifying entorhinal input to the DG during behaviorally relevant states, such a social memory encoding, enabling coordinated regulation of excitability and synaptic strength across the circuit.

### Summary

In summary, our findings suggest that the dentate gyrus may be a site of oxytocin action within the hippocampal formation. By enhancing both intrinsic excitability and synaptic drive, oxytocin increases the responsiveness of granule cells to perforant path input and engages mechanisms overlapping with activity-dependent plasticity. We suggest that oxytocin may prime DG GCs for synaptic strengthening during salient experiences. This dual modulation of excitability and synaptic efficacy extends the cellular actions of oxytocin to include DG GCs, suggesting that oxytocin acts at multiple levels, both presynaptic and postsynaptic, to coordinate hippocampal computation and plasticity, providing a cellular framework through which oxytocin could promote the encoding of socially and behaviorally relevant stimuli. More broadly, these results expand the known influence of oxytocin from higher-order social circuits to fundamental hippocampal computations, highlighting a distributed oxytocinergic mechanism that links social signaling with memory circuit plasticity.

## Additional Information

Competing interests: No competing interests declared

## Author contributions

A.M.M. and D.H.B. conceptualized and designed the research. A.M.M. and B.B. performed all the experiments. A.M.M. and B.B. analyzed the data. A.M.M. and D.H.B. interpreted the results of the experiments. A.M.M., B.B., and D.H.B. drafted the manuscript. A.M.M. All persons designated as authors qualify for authorship and all those who qualify for authorship are listed.

## Data Availability Statement

The data that support the findings of this study are available from the corresponding author upon reasonable request.

## Funding

R01 MH13317 (DHB) 1T23HL456789-38 (AMM)

## Notes

### Competing Interest Statement

The authors have declared no competing interest.

### Summary of Updates

Author list updated to correct order, with Dr. Darrin Brager listed as senior author.

